## Supplemental figures for "Plant SYP12 syntaxins mediate an evolutionarily conserved general immunity to filamentous pathogens"

### Figure S1

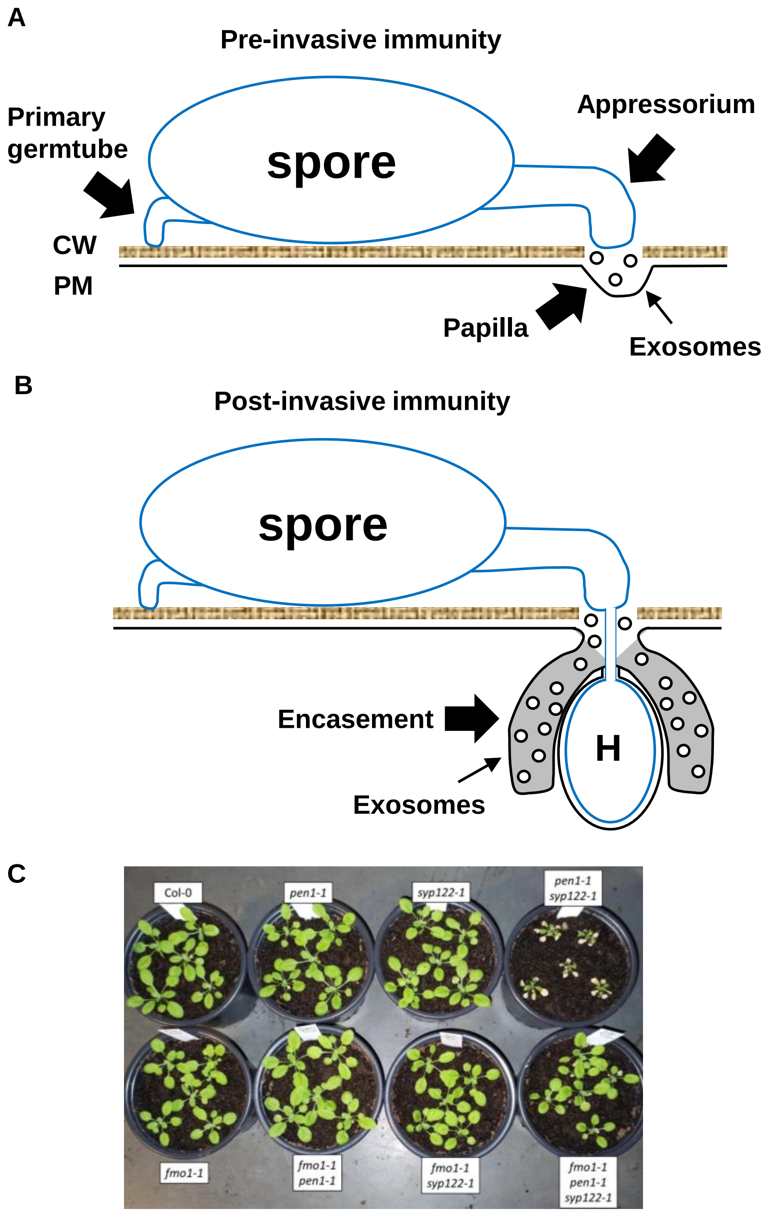


**Fig. S1. Pre- and post-invasive immunity.** (A) Attack by non-adapted filamentous pathogens such as *Bgh* on *Arabidopsis*, is met by a localized apposition (papilla) formed between the plant cell wall (CW) and the plasma membrane (PM), containing callose, phenolic compounds, reactive oxygen species, and exosomes, that likely prevents penetration (Zeyen et al., 2002; Assaad et al., 2004; An et al., 2006). (B) Upon successful penetration, the host cell forms an encasement, similar in composition to the papilla, that eventually encloses the developing haustorium (H) and prevents nutrient uptake. Similar to *Bgh*, spores of *C. destructivum* and *P. infestans* attempt to penetrate and form an intracellular pathogenic structure (IPS, haustoria, biotrophic hypae and infection vesicle, respectively). In the case of an ineffective encasement response, the host cell initiates a cell death response that prevents pathogen growth. (C) Plants at ~4 weeks, showing the growth phenotypes of the mutant lines used in these studies.

### Figure S2

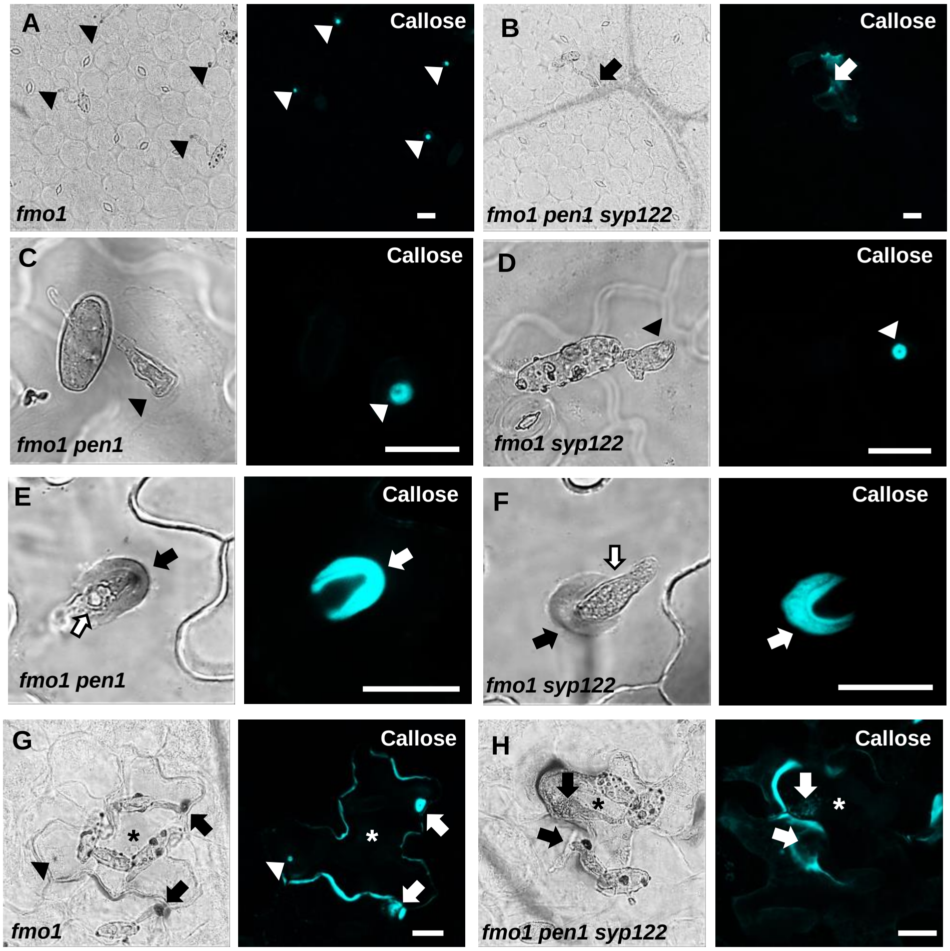


**Fig. S2. PEN1/SYP122 are required for papilla and encasement responses to *Bgh*.** (A-H) Accumulation of callose in response to *Bgh* attack at (A-D) non-penetrated (arrowheads) and (E-H) penetrated (arrows) attack sites. Open arrows point to the developing IPS. * marks cells with cell death response. Bars = 20 µm.

### Figure S3

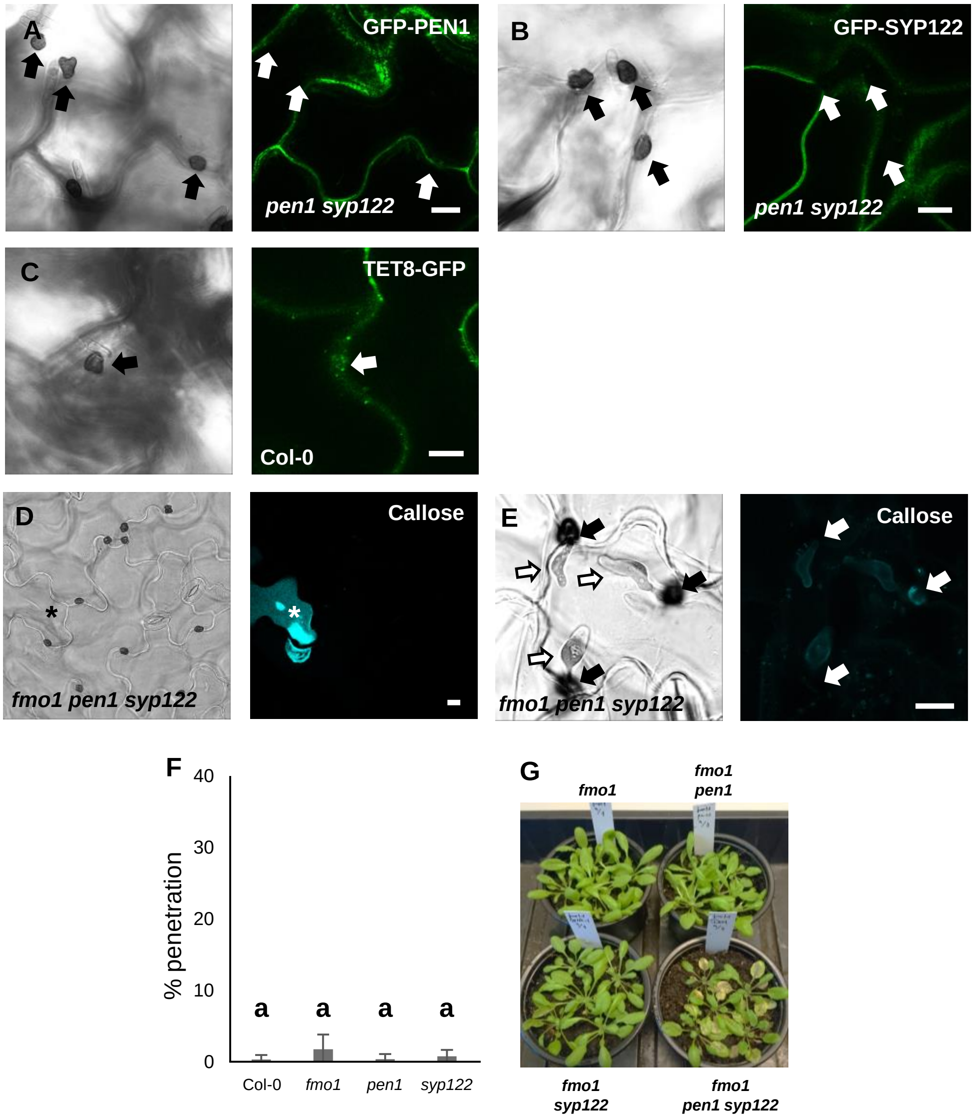


**Fig. S3. PEN1/SYP122 are required for pre-invasive immunity towards *C. destructivum*.** (A-C) Localisation of GFP-PEN1 (A), GFP-SYP122 (B) and TET8-GFP (C) in response to attack by *C. destructivum*. (D-E) Accumulation of callose in response to attack by *C. destructivum* in (D) non-penetrated cells and (E) at penetration sites (arrows). Open arrows point to the developing IPS. * marks cell with cell death response. (F) Frequency of penetration by *C. destructivum*. (G) Disease symptoms at 5 dai with *C. destructivum*. Bars = 10 µm. (F) All values are mean ±SD (n = 4 leaves per genotype). Different letters indicate significantly different values at P ≤ 0.001 estimated using logistic regression.

### Figure S4

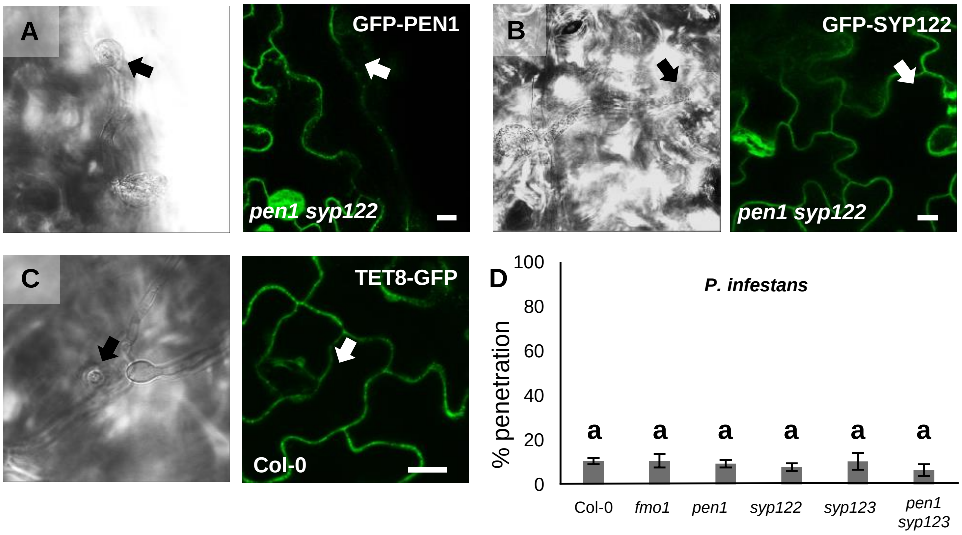


**Fig. S4. PEN1/SYP122 are required for pre-invasive immunity towards *P. infestans*.** (A-C) Localisation of GFP-PEN1 (A), GFP-SYP122 (B) and TET8-GFP (C) in response to attack by *P. infestans*. (D) Frequency of penetration by *P.infestans*. Bars = 10 µm. (D) All values are mean ±SD (n = 4 leaves per genotype). Different letters indicate significantly different values at P ≤ 0.001 estimated using logistic regression.

### Figure S5

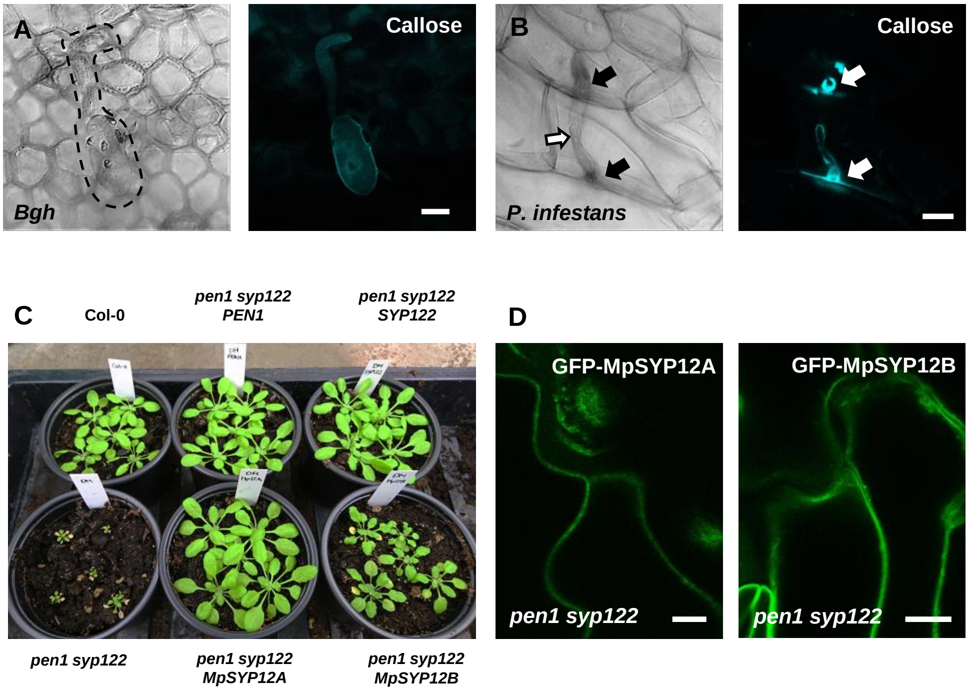


**Fig. S5. Responses in *Marchantia polymorpha* to filamentous pathogens.** (A) On *M. polymorpha*, spores from *Bgh* (dotted line) were mis-differentiated with no or wrongly orientated appressoria that did not attack the host-cell. (B) Accumulation of callose in *M. polymorpha* in response to attack by *P. infestans* (arrows) in penetrated cells. Open arrow points to the developing IPS. Note that the frequency of *P. infestans* spores that attack a host-cell is very low. Bars=10 µm.

### Figure S6

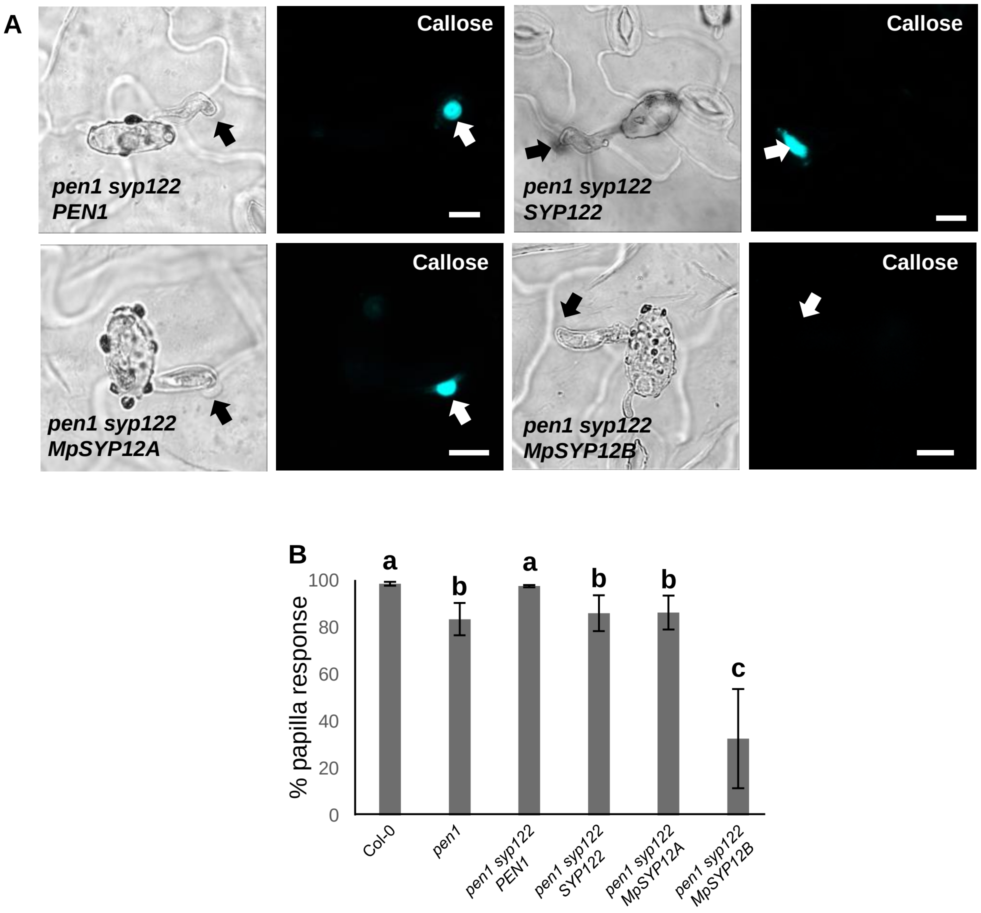


**Fig. S6. *Marchantia* syntaxins rescue papilla responses.** (A) Accumulation of callose in response to *Bgh* attack at non-penetrated (arrows) attack sites. Bars = 10 µm. (B) Frequency of papillae in response to *Bgh* in non-penetrated cells. All values are mean ±SD (n = 5 leaves per genotype). Different letters indicate significantly different values at P ≤ 0.001 estimated using logistic regression.

### Figure S7

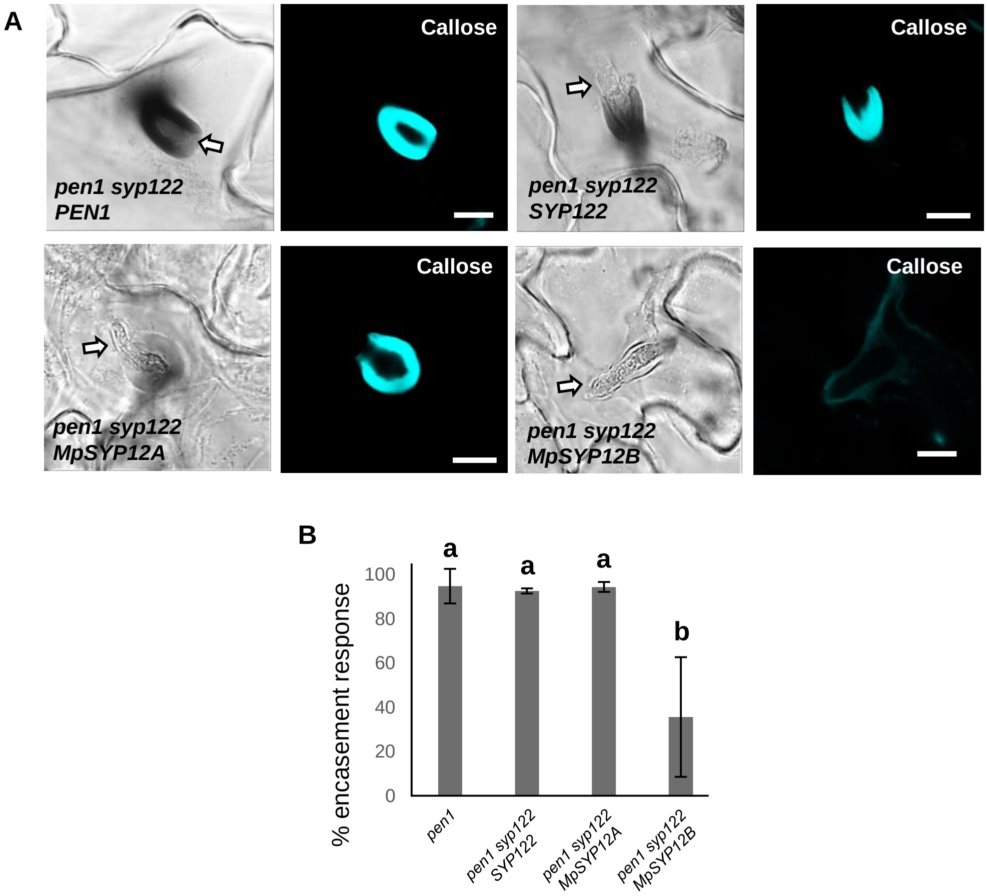


**Fig. S7. *Marchantia* syntaxins rescue encasement responses.** (A) Accumulation of callose in response to *Bgh* attack at successful penetration sites. Open arrows point to the developing IPS. Bars = 10 µm. (B) Frequency of encasements in response to *Bgh* haustoria in penetrated cells. All values are mean ±SD (n = 5 leaves per genotype). Different letters indicate significantly different values at $P \leq 0.001$ estimated using logistic regression.

### Figure S8

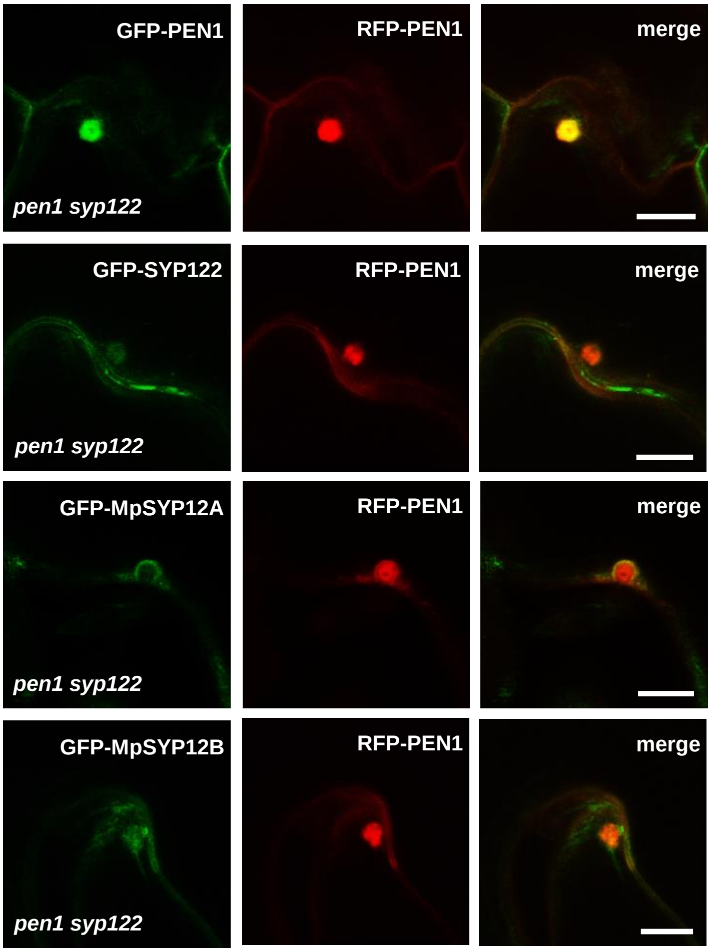


**Fig. S8. Localization of SYP12 clade members at *Bgh* attack sites.** Comparative papilla localization of SYP12 members from *Arabidopsis* or *Marchantia* in response to attack by *Bgh* in non-penetrated cells. Bars = 10 µm.

### Figure S9

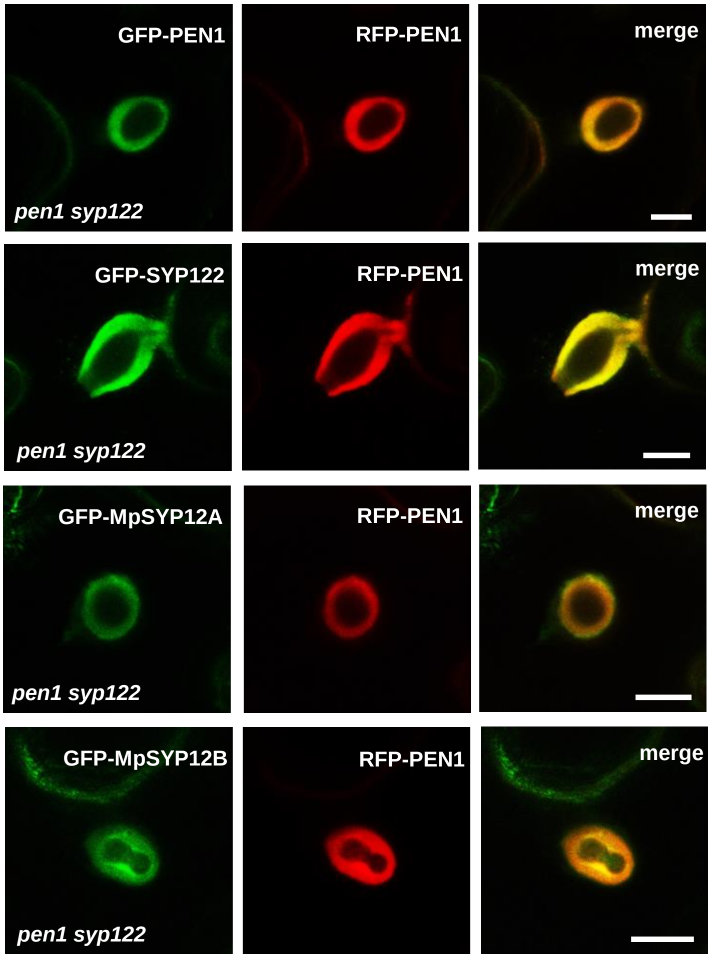


**Fig. S9. Localization of SYP12 clade members at *Bgh* penetration sites.** Comparative encasement localization of SYP12 members from *Arabidopsis* or *Marchantia* in response to haustoria by *Bgh* in penetrated cells. Bars = 10 µm.

### Figure S10

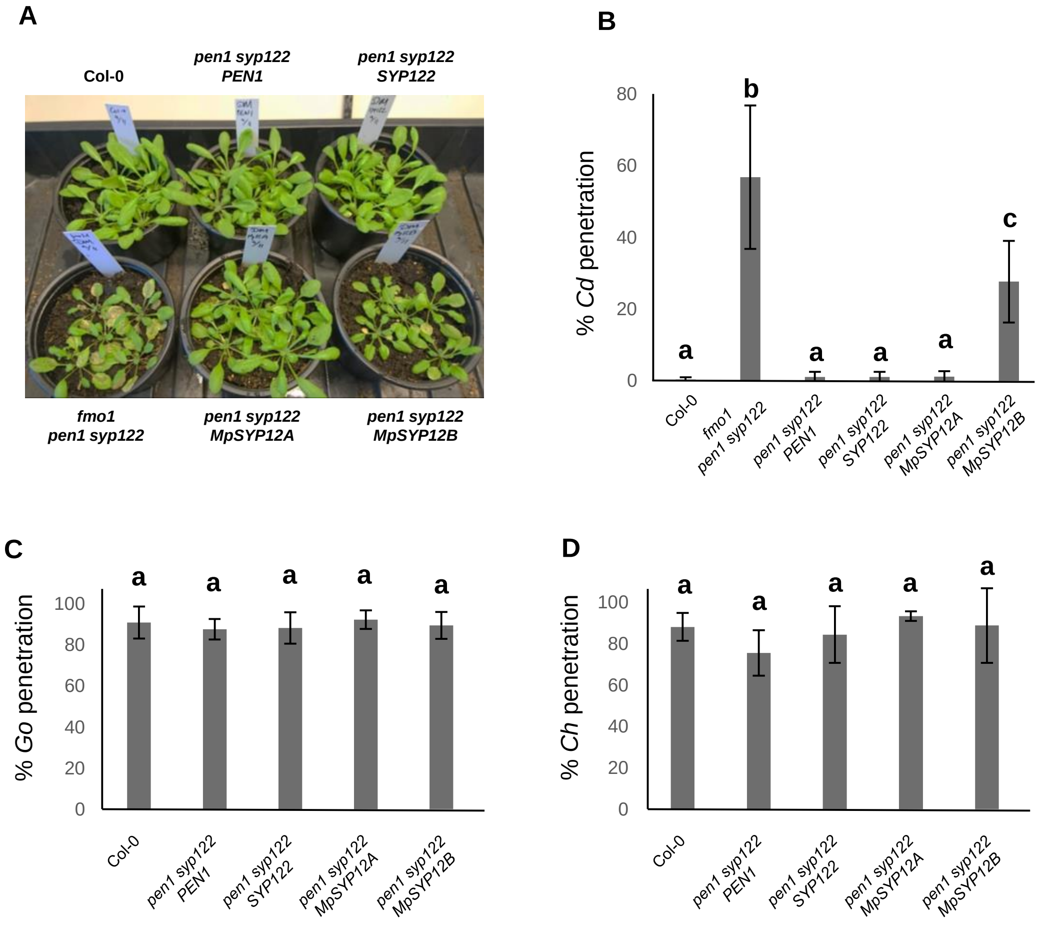


**Fig. S10. *Marchantia* SYP12s restore immunity in *Arabidopsis* towards *C. destructivum*.** (A) Disease symptoms at 5 dai with *C. destructivum*. (B-D) Frequency of penetration by *C. destructivum* (B), *G. orontii* (C) and *C. higginsianum* (D). (B-D) All values are mean ±SD (n = 4 leaves per genotype). Different letters indicate significantly different values at P ≤ 0.001 estimated using logistic regression.

### Figure S11

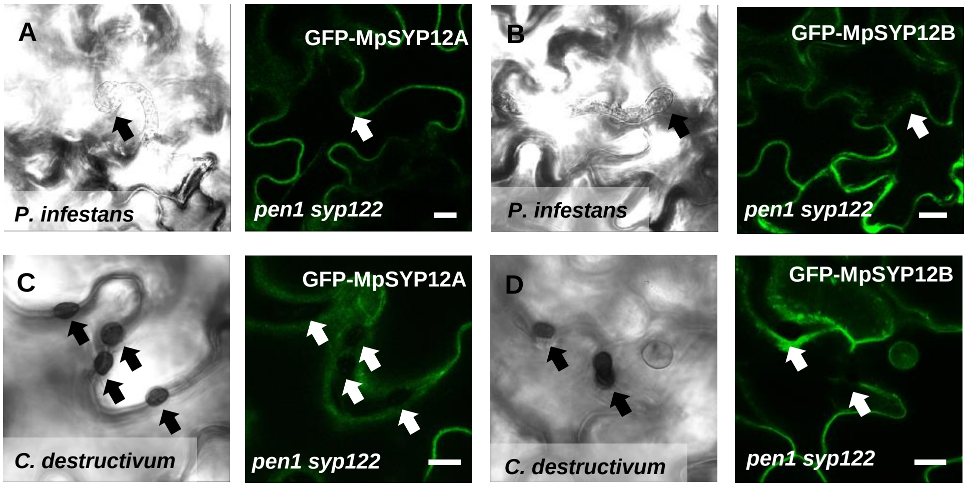


**Fig. S11. Localization of *Marchantia* SYP12 clade members at attack sites.** (A-D) Localization of GFP-MpSYP12A (A, C) and GFP-MpSYP12A (B, D) in response to attack by *P. infestans* (A-B) and *C. destructivum* (C-D) in non-penetrated cells. Bars = 10 µm.
